# The Genetic Landscape of Diamond-Blackfan Anemia

**DOI:** 10.1101/365890

**Authors:** Jacob C. Ulirsch, Jeffrey M. Verboon, Shideh Kazerounian, Michael H. Guo, Daniel Yuan, Leif S. Ludwig, Robert E. Handsaker, Nour J. Abdulhay, Claudia Fiorini, Giulio Genovese, Elaine T. Lim, Aaron Cheng, Beryl B. Cummings, Katherine R. Chao, Alan H. Beggs, Casie A. Genetti, Colin A. Sieff, Peter E. Newburger, Edyta Niewiadomska, Michal Matysiak, Adrianna Vlachos, Jeffrey M. Lipton, Eva Atsidaftos, Bertil Glader, Anupama Narla, Pierre-Emmanuel Gleizes, Marie-Françoise O’Donohue, Nathalie Montel-Lehry, David J. Amor, Steven A. McCarroll, Anne H. O’Donnell-Luria, Namrata Gupta, Stacey B. Gabriel, Daniel G. MacArthur, Eric S. Lander, Monkol Lek, Lydie Da Costa, David. G. Nathan, Andrei K. Korostelev, Ron Do, Vijay G. Sankaran, Hanna T. Gazda

## Abstract

Diamond-Blackfan anemia (DBA) is a rare bone marrow failure disorder that affects 1 in 100,000 to 200,000 live births and has been associated with mutations in components of the ribosome. In order to characterize the genetic landscape of this genetically heterogeneous disorder, we recruited a cohort of 472 individuals with a clinical diagnosis of DBA and performed whole exome sequencing (WES). Overall, we identified rare and predicted damaging mutations in likely causal genes for 78% of individuals. The majority of mutations were singletons, absent from population databases, predicted to cause loss of function, and in one of 19 previously reported genes encoding for a diverse set of ribosomal proteins (RPs). Using WES exon coverage estimates, we were able to identify and validate 31 deletions in DBA associated genes. We also observed an enrichment for extended splice site mutations and validated the diverse effects of these mutations using RNA sequencing in patientderived cell lines. Leveraging the size of our cohort, we observed several robust genotype-phenotype associations with congenital abnormalities and treatment outcomes. In addition to comprehensively identifying mutations in known genes, we further identified rare mutations in 7 previously unreported RP genes that may cause DBA. We also identified several distinct disorders that appear to phenocopy DBA, including 9 individuals with biallelic *CECR1* mutations that result in deficiency of ADA2. However, no new genes were identified at exome-wide significance, suggesting that there are no unidentified genes containing mutations readily identified by WES that explain > 5% of DBA cases. Overall, this comprehensive report should not only inform clinical practice for DBA patients, but also the design and analysis of future rare variant studies for heterogeneous Mendelian disorders.

## INTRODUCTION

Diamond-Blackfan anemia (DBA), originally termed congenital hypoplastic anemia, is an inherited bone marrow failure syndrome estimated to occur in 1 out of 100,000 to 200,000 live births^1; 2^. A consensus clinical diagnosis for DBA suggests that individuals with this disorder should be diagnosed within the first year of life and have normochromic macrocytic anemia, limited cytopenias of other lineages, reticulocytopenia, and a visible paucity of erythroid precursor cells in the bone marrow^3^. Nonetheless, an increasing number of cases that fall outside of these strict clinical criteria are being recognized^4^. Treatment with corticosteroids can improve the anemia in 80% of cases, but individuals often become intolerant to long-term corticosteroid therapy and turn to regular red blood cell transfusions, the only available standard therapy for the anemia^5^. Currently, a hematopoietic stem cell transplant (HSCT) is the sole curative option, but this procedure carries significant morbidity and is generally restricted to those with a matched related donor^6^. Ultimately, 40% of cases remain dependent upon corticosteroids which increase the risk of heart disease, osteoporosis, and severe infections, while another 40% become dependent upon red cell transfusions which requires regular chelation to prevent iron overload and increases the risk of alloimmunization and transfusion reactions, both of which can be severe co-morbidities^2;5^.

In contrast to many other rare, presumed monogenic or Mendelian disorders^7-9^, putative causal genetic lesions have now been identified in an estimated 50-60% of DBA cases^2^. In 1999, mutations in ribosomal protein S19 (RPS19), one of the proteins in the 40S small ribosomal subunit, were identified as the first causal genetic lesions for DBA in ~25% of cases^10^. Through the use of targeted Sanger sequencing, whole exome sequencing (WES), and copy number variant (CNV) assays, putatively causal haploinsufficient mutations have been identified in 19 of the 79 ribosomal protein (RP) genes (RPS19, RPL5, RPS26, RPL11, RPL35A, RPS10, RPS24, RPS17, RPS7, RPL26, RPL15, RPS29, RPS28, RPL31, RPS27, RPL27, RPL35, RPL18, RPS15A) making DBA one of the best genetically defined congenital disorders. In 2012, through the use of unbiased WES, mutations in *GATA1*, a hematopoietic master transcription factor that is both necessary for proper erythropoiesis and sufficient to reprogram alternative hematopoietic lineages to an erythroid fate, were identified as the first non-RP mutations in DBA^11; 12^. Further studies on GATA1 and other novel genes mutated in DBA, including the RPS26 chaperone protein TSR2^13; 14^, have provided new insights into the pathogenesis of this disorder, suggesting that DBA results from impaired translation of key erythroid transcripts, such as the mRNA encoding *GATA1*, in early hematopoietic progenitors which ultimately impairs erythroid lineage commitment^14-18^.

Given the success of unbiased WES in identifying pathogenic mutations in many Mendelian disorders^7; 8; 11; 13; 19; 20^, we recruited and performed sequencing on a large cohort of 472 affected individuals, the size of which is equivalent to 6 years of spontaneous DBA births in the USA, Canada, and Europe, containing individuals with a clinical diagnosis or strong suspicion of DBA. In this report, we describe the results of an exhaustive genetic analysis of this cohort and discuss our experience of attempting to achieve comprehensive molecular diagnoses, while limiting false positive reports.

## MATERIALS AND METHODS

### Whole Exome Sequencing

A total of 445 affected individuals and 72 unaffected family members underwent whole exome sequencing at the Broad Institute (dbGAP accession phs000474.v2.p1). Generally, whole exome sequencing and variant calling was performed as previously reported with several modifications^11^. Library construction was performed as described in Fisher et al.^21^, with the following modifications: initial genomic DNA input into shearing was reduced from 3 μg to 10-100 ng in 50 μL of solution. For adapter ligation, Illumina paired end adapters were replaced with palindromic forked adapters, purchased from Integrated DNA Technologies, with unique 8 base molecular barcode sequences included in the adapter sequence to facilitate downstream pooling. With the exception of the palindromic forked adapters, the reagents used for end repair, A-base addition, adapter ligation, and library enrichment PCR were purchased from KAPA Biosciences in 96-reaction kits. In addition, during the post-enrichment SPRI cleanup, elution volume was reduced to 20 μL to maximize library concentration, and a vortexing step was added to maximize the amount of template eluted.

For Agilent capture, in-solution hybrid selection was performed as described by Fisher et al.^21^, with the following exception: prior to hybridization, two normalized libraries were pooled together, yielding the same total volume and concentration specified in the publication. Following post-capture enrichment, libraries were quantified using quantitative PCR (kit purchased from KAPA Biosystems) with probes specific to the ends of the adapters. This assay was automated using Agilent’s Bravo liquid handling platform. Based on qPCR quantification, libraries were normalized to 2 nM and pooled by equal volume using the Hamilton Starlet. Pools were then denatured using 0.1 N NaOH. Finally, denatured samples were diluted into strip tubes using the Hamilton Starlet.

For ICE capture, in-solution hybridization and capture were performed using the relevant components of Illumina’s Rapid Capture Exome Kit and following the manufacturer’s suggested protocol, with the following exceptions: first, all libraries within a library construction plate were pooled prior to hybridization. Second, the Midi plate from Illumina’s Rapid Capture Exome Kit was replaced with a skirted PCR plate to facilitate automation. All hybridization and capture steps were automated on the Agilent Bravo liquid handling system. After post-capture enrichment, library pools were quantified using qPCR (automated assay on the Agilent Bravo), using a kit purchased from KAPA Biosystems with probes specific to the ends of the adapters. Based on qPCR quantification, libraries were normalized to 2 nM, then denatured using 0.1 N NaOH on the Hamilton Starlet. After denaturation, libraries were diluted to 20 pM using hybridization buffer purchased from Illumina.

Cluster amplification of denatured templates was performed according to the manufacturer’s protocol (Illumina) using HiSeq v3 cluster chemistry and HiSeq 2000 or 2500 flowcells. Flowcells were sequenced on HiSeq 2000 or 2500 using v3 Sequencingby-Synthesis chemistry, then analyzed using RTA v.1.12.4.2 or later. Each pool of whole exome libraries was run on paired 76 bp runs, with an 8 base index sequencing read was performed to read molecular indices

### Variant Calling and Annotation

We performed joint variant calling for single nucleotide variants and indels across all samples in this cohort and ~6,500 control samples from the Exome Sequencing Project (http://evs.gs.washington.edu/EVS/) using GATK v3.4. Specifically, we used the HaplotypeCaller pipeline according to GATK best practices (https://software.broadinstitute.org/gatk/). Variant quality score recalibration (VQSR) was performed, and in the majority of analyses only “PASS” variants were investigated. The resultant variant call file (VCF) was annotated with Variant Effect Predictor v91^22^, Loftee, dpNSFP-2.9.3^23^, and MPC^24^. A combination of GATK^25^, bcftools, and Gemini^26^ was used to identify rare and predicted damaging variants. Specifically, variants with an allele count (AC) of ≤ 3 in gnomAD (a population cohort of 123,136 exomes) were considered rare, and variants annotated as LoF (splice acceptor or donor variants, stop gained, stop lost, start lost, and frameshifts) or missense by VEP were considered potentially damaging. Other rare variants in previously described DBA genes with other annotations or no annotation were investigated on a case by case basis. When family members had also undergone WES, variants were required to fit Mendelian inheritance (e.g. dominant for RP genes, hemizygous for *GATA1* and *TSR2*). In each family, all rare and predicted damaging *de novo* or recessive mutations were also considered. In all cases, pathogenic variants reported by Clinvar, as well as rare variants in genes known to cause other disorders of red cell production or bone marrow failure were also considered^27^. All putative causal variants were manually inspected in IGV^28^. Cohort quality control including the ancestry analysis, crypic relatedness and sex checks was performed using peddy^29^. Specifically, PCA was performed on 1000 Genomes project samples for the overlap of variants measured in the DBA cohort with ≈ 25,000 variants from samples in the 1000 Genomes project. DBA cohort samples were then projected onto these PCs, and ancestry in the DBA cohort was predicted from the PC coordinates using a support vector machine trained on known ancestry labels from 1000 Genomes samples. Relatedness parameters were calculated (coefficient of relatedness, ibs0, ibs1, ibs2) using these variants and were compared to known relationships from the cohort pedigrees; cases which did not agree were manually validated and corrected. In all cases, sex checks (presence of heterozygous variants on the X chromosome) performed by peddy aligned with available cohort information.

### Targeted Sanger Sequencing

The Primer3 program (http://frodo.wi.mit.edu/primer3/) was used to design primers to amplify a fragment of 200–300 bp targeting a specific region of either exon or intron of the gene of interest. Polymerase chain reaction was performed using Dream Taq Polymerase (Life Science Technology, Cat# EP0701) and 30 μg of genomic DNA in a 15 μl reaction. The reaction was performed with an initial denaturation of 5 min at 94°C followed by 29 cycles of second denaturation at 94°C for 45 sec, annealing at 57°C for 45 sec, and extension at 72°C for 45 sec. The final extension was performed at 72°C for 10 min. The PCR product was treated with the reagent ExoSAP-IT (USB, Santa Clara, CA) and submitted for Sanger sequencing to the Boston Children’s Hospital Molecular Genetics Core Facility. The resulted sequences were analyzed using Sequencher 4.8 software (Gene Codes, Ann Arbor, MI) and compared with normal gene sequence provided through the UCSC Genome Browser.

### Lymphoblastoid Cell Lines

To generate lymphoblastoid cell lines from peripheral blood, Histoplaque solution was used to isolate the buffy coat containing mononuclear cells. Mononuclear cells were transferred into a new tube and washed twice with PBS. Cells were resuspended into 2 ml complete RPMI 1640 containing 15% FBS and 5% Penicillin/Streptomycin and Glutamine. 2 ml of Epstein-Barr virus (EBV) solution was added, and cells were incubated at 37°C and 5% CO_2_ overnight. After adding 5 ml complete RPMI, cells were allowed to grow to confluency and maintained using the regular cell culture procedure. Epstein-Barr virus (EBV) was generated by growing B95-8 cells in RPMI complete until they were at a high cell concentration (1™2×10^9^) for 12 to 14 days. Cells were centrifuged at 1,300 RPM for 10 minutes at 20°C. The supernatant (containing EBV virus) was passed through a 0.45 μm PEB filter twice, aliquoted in 2 ml cryogenic vials, and stored at −80°C. This procedure was performed in accordance with the Boston Children’s Hospital’s Biosafety protocol.

### RNA-seq and Splicing Analysis

RNA was isolated using RNeasy kits (Qiagen) according to the manufacturer’s instructions. 1-20 ng of RNA were forwarded to a modified Smart-seq2 protocol and after reverse transcription, 8-9 cycles of PCR were used to amplify transcriptome libraries^30^. Quality of whole transcriptome libraries were validated using a High Sensitivity DNA Chip run on a Bioanalyzer 2100 system (Agilent), followed by sequencing library preparation using the Nextera XT kit (Illumina) and custom index primers. Sequencing libraries were quantified using a Qubit dsDNA HS Assay kit (Invitrogen) and a High Sensitivity DNA chip run on a Bioanalyzer 2100 system (Agilent). All libraries were sequenced using Nextseq High Output Cartridge kits and a Nextseq 500 sequencer (Illumina). Libraries were sequenced paired-end (2 × 38 cycles).

Fastq files were aligned to the Ensembl GRCh37 r75 genome assembly (hg19) using 2-Pass STAR alignment^31; 32^. Based on the general approach previously described in Cummings et al.^33^, STAR first pass parameters were adjusted as follows in order to more inclusively detect novel splice junctions: -“-outSJfilterCountTotalMin 10 10 10 10 --outSJfilterCountUniqueMin −1 −1 −1 −1 --alignIntronMin 20 --alignIntronMax 1000000 --alignMatesGapMax 1000000 --alignSJoverhangMin 8 --alignSJDBoverhangMin 3 --outSJfilterOverhangMin 0 0 0 0 --outSJfilterDistToOtherSJmin 0 0 0 0 --scoreGenomicLengthLog2scale 0”. Novel junctions detected in the first pass alignment were combined and included as candidate junctions in the second pass. Candidate genes were investigated for splicing using both IGV^28^ and the Gviz package^34^. Sashimi plots were created using Gviz. Gene expression was quantified using RSEM^35^, and expression differences were determined by the log^2^ fold change in transcripts per million (TPM).

### Copy Number Variant Identification and Validation

Copy number variant (CNV) analysis was performed for the entire cohort using XHMM separately for ICE and Agilent exomes, as previously described^36; 37^. Specifically, XHMM takes as input a sample by exon read coverage matrix, performs principal component (PC) analysis, re-projects the matrix after removing PCs that explain a large proportion of the variance, normalizes the matrix (z-score), then uses a hidden Markov model (HMM) to estimate copy number state. For known RP genes, candidate deletions were nominated either by (1) XHMM deletion calls or (2) manual investigation of outliers in the z-score distribution for each exon. When WES was performed in other family members, the inheritance of putative CNVs was also determined. Putative CNVs were validated using ddPCR^38^. Specifically, primers and probes were designed to amplify exons with putative deletions. 50 ng of DNA per sample (at least one test and one control per reaction) were digested with a restriction enzyme, either Hind or HaeIII, and master mixes containing FAM targeted assays and control HEX RPP30 assays. Subsequently, plates were foil sealed, vortexed and placed in an autodroplet generator (BioRad). Once the droplets were generated, plates were placed in thermal cycler C1000 Touch (BioRad) for DNA amplification. PCR was performed with an initial denaturing step at 95°C for 10 min, followed by 40 cycles of denaturing at 95°C for 30 sec and annealing at 60°C for 1 min. Subsequently, enzyme deactivation was achieved by heating to 98°C for 10 min. Each PCR run included no-template controls and normal controls. The results of ddPCR were generated using QX200 Droplet Reader (BioRad) and analyzed using QuantaSoft Analysis Pro (BioRad).

### Segmental Duplication Analysis

To investigate the copy number distribution of *RPS17* in the human population, we used Genome STRiP^39^ to determine the copy number of this gene using whole genome sequence data from the 1000 Genomes Project^40^ in 2,535 individuals of diverse ancestry. We first measured the copy number of the segmental duplication containing *RPS17*, specifying the coordinates of both copies of the segmental duplication (hg19 coordinates chr15:82629052-82829645 and chr15:83005382-83213987) to estimate the total copy number (which would be 4 for individuals homozygous for the hg19 reference haplotype that contains two copies of this segment). We further measured just a 5 kb segment directly at *RPS17* (hg19 coordinates chr15:83205001-83210000 and chr15:82820658-82825658) to determine whether the gene itself was present in two diploid copies in individuals from the 1000 Genomes cohort. We also performed the same measurement on a control locus, a true segmental duplication of similar size (approximately 200 kb) on chromosome 5 which appears to exhibit no copy number variation in the 1000 Genomes cohort (hg19 coordinates chr5:175350365-175558672 and chr5:177133499-177347466).

### Penetrance Analysis

Penetrance analysis was performed as previously described in Minikel et al.^41^ with a few key modifications. Specifically, we use Bayes’ rule to obtain *P*(*D*|*G*) = *P*(*D*) × *P*(*G*|*D*)*/P*(*G*), where *P*(*D*|*G*) is the penetrance for a specific genotype *G*, *P*(*D*) is the lifetime risk of the disease *D* in a general population, *P*(*G*|*D*) is the proportion of individuals with DBA who have the specific genotype *G*, and *P*(*G*) is the proportion of individuals in the general population who have the specific genotype. We can calculate *P*(*D*) as *average lifetime* × *DBA incidence* = 80 *years* × 7/1000000. We obtain an estimate for *P*(*G*|*D*) as # *individuals with* ≥ 1 *allele in DBA cohort*. Similarly, we can obtain an estimate for *P*(*G*) as # *individuals with* ≥ 1 *allele in gnomAD* + 1, where we add 1 (estimated integer of DBA cases in a population of size 121,136) to the proportion of individuals with the specific genotype in gnomAD, since gnomAD is not perfectly representative of the general population and most or all potential DBA cases are likely to have been removed. This allows us to plugin to calculate a point estimate for *P*(*D*|*G*) as min(1,*P*(*D*) × *P*(*G*|*D*)/*P*(*G*)). Similarly, we can quantify the spread in this estimate using 95% Wilson confidence intervals of a binomial distribution (also known as score intervals). We note that by adding 1 to the denominator that this could potentially result in a slightly lower and more conservative estimate of penetrance. Since the majority of variants identified were singletons and we are primarily interested in inference at the gene and variant type level, we collapsed variants by predicted effects (LoF, missense) and gene in order to obtain more robust estimates. A max total allele count (AC) of 12 across the combined set of DBA and gnomAD exomes was used as a filter, since a few variants reached higher prevalence in DBA. The penetrance of the mutation identified as polymorphic from the DBAgenes database (RPL5:c.418G>A) was estimated using the same formula, and *P*(*G*|*D*) was estimated using the DBA cohort in this study (1 individual was observed to have the A allele).

### Structural Analysis

The cryo-EM structure of the human 80S ribosome^42^ and P-stalk proteins from the cryo-EM structure of the yeast 80S ribosome^43^ were used to create a hybrid 80S structural model shown in Figure 1E. Structural superposition, analyses, and figures were rendered using PyMOL^44^.

**Figure 1.**
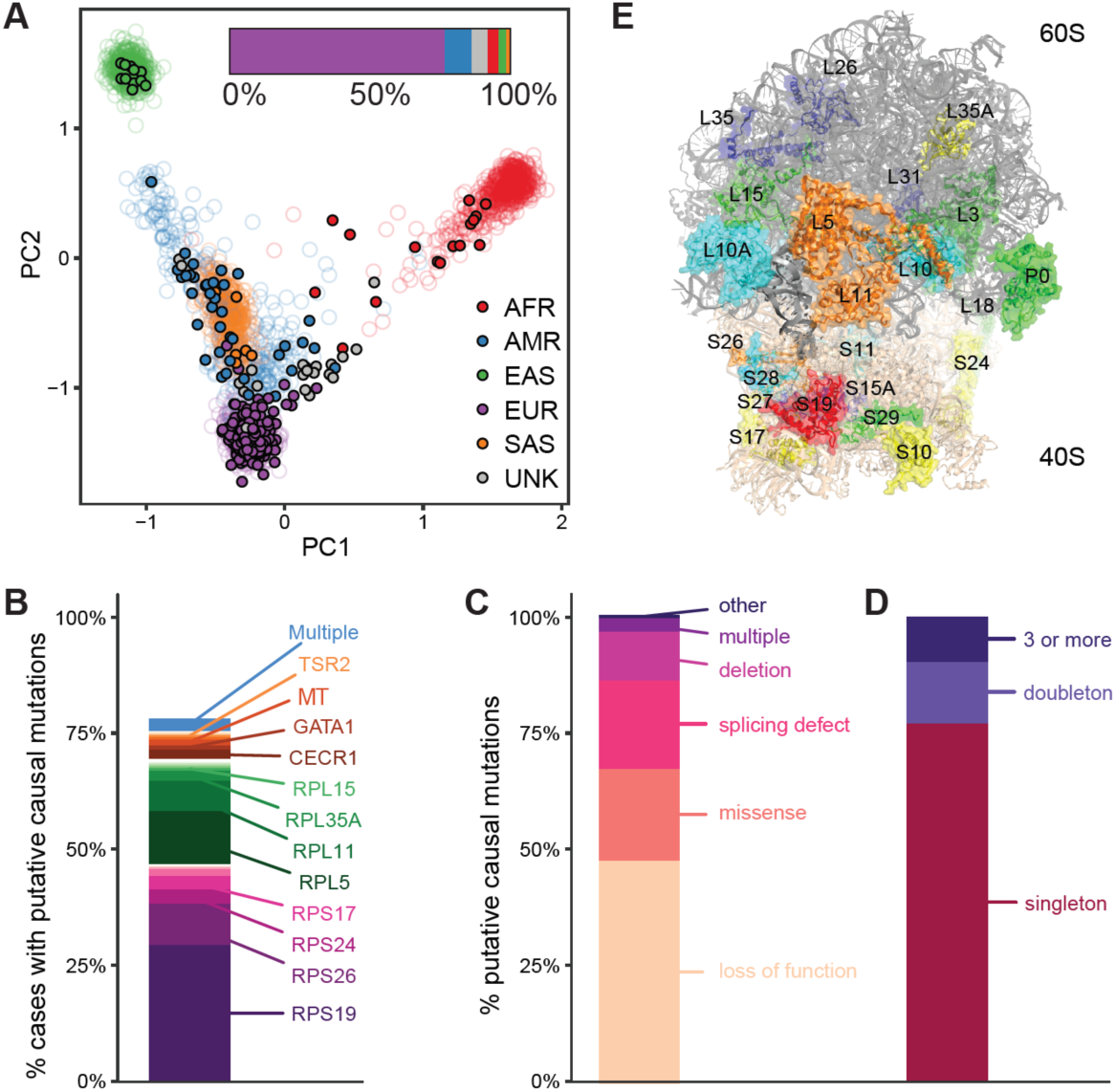
Mutational Spectrum of Likely Pathogenic Variants in DBA. (**A**) PCA of genetic ancestry based upon 1000 Genomes for the DBA cohort. Filled circles represent individual DBA families and open circles represent 1000 Genomes individuals. (**B**) Percentage of putative causal mutations in each gene. A total of 78% of cases have a putative causal mutation. (**C**) Types of putative causal mutations. (**D**) Relative frequency of putative causal mutations in the DBA cohort. (**E**) Structure of the assembled ribosome highlighting the 19 known and 7 novel RP genes that are mutated in the DBA cohort. The color coding reflects the frequency of mutations: blue (none found in this study; but reported in other studies), cyan (1 mutation), green (2-3 mutations), yellow (7-15 mutations), orange (31-54 mutations) and red (more than 100 mutations). RPS7, RPL27, RPL34, and RPL19 are mutated in DBA but are obscured by other proteins from this viewpoint.

### Gene Burden Analyses

Gene-based burden testing^45^ was performed using a one-sided Fisher’s exact test of the 2×2 table of genotype counts (genotype present or genotype absent) between cases (407 unrelated cases from the DBA cohort) and controls (gnomAD). gnomAD is an aggregation database of exome sequencing from 123,136 individuals who are not known to have a severe Mendelian condition (http://gnomad.broadinstitute.org)^46^. Counts under the dominant model were generated for DBA by counting the number of individuals who carry at least one qualifying variant in each gene and for gnomAD by summing the allele counts for qualifying variants in each gene. For the recessive model, counts in DBA were generated by counting the number of individuals who carry two or more qualifying variants in a gene, or who are homozygous for a qualifying variant. For the recessive model in gnomAD, the number of individuals who carry a homozygous variant was added to a predicted number of compound heterozygous variant carriers. The predicted number of compound heterozygous variant carriers was calculated by squaring the total heterozygous variant carrier rate in each gene and multiplying by the total sample size. P-values < 2.5 × 10^−6^ were considered significant (of 0.05 corrected for testing ≈ 20,000 genes). Predicted damaging missense mutations were identified using PolyPhen2^47^. Several steps were taken to match variant call set quality since variants were not jointly called for the cases and controls. First, read depth was computed in each cohort separately and only sites where the read depth was > 10 in each cohort were included. Second, sites present in low complexity regions were removed. Third, rare synonymous variant burden testing was performed for different variant quality score recalibration (VQSR) combinations until the -log^10^ p-values from the Fisher’s exact test followed the expected distribution. Specifically, inclusion of the top 85% of VQSR variants from the DBA cohort and the top 95% of VQSR variants from the gnomAD cohort resulted in the best fit (erring slightly to be more conservative than not).

### Statistical Analyses

In order to test for differences in outcome (e.g. congenital abnormalities, treatment outcomes), a Fisher’s exact test was performed on the genotype by phenotype count matrix and p-values were calculated from 100,000 Monte Carlo simulations. The type of mutation (e.g. LoF or missense) was not separately investigated, since this is confounded by the exact gene implicated, although gene-phenotype associations were secondarily validated after removing missense mutations. For outcomes of interest, 95% binomial confidence intervals are reported in addition to the point estimates. Precision recall curves for classification of RPS19 missense mutations as belonging to the DBA cohort or to the gnomAD cohort based upon missense pathogenicity predictive methods were calculated using the R package pROC. Other enrichment tests (e.g. splicing position) were calculated using Fisher’s exact tests on 2×2 tables of variant counts. Power analysis for burden testing was performed using the power.fisher.test function in the R statmod package with at least 10,000 simulations.

## RESULTS

### Rare Loss of Function and Missense Variants in Known DBA Genes

We assembled a cohort of 472 individuals with a likely diagnosis of DBA of predominantly European descent (76%) from DBA registries and clinicians over the course of 20 years and performed WES on 94% in an attempt to comprehensively identify or verify causal mutations (**Fig. 1A**). Combining all approaches (**Materials and Methods**), we identified putative causal mutations for the observed anemia in 78% (369/472) of cases (**Fig. 1B**). The majority of these mutations were in one of the 19 previously known DBA genes (330/472, 70%) and were primarily rare (gnomAD AC ≤ 3) loss of function (LoF) or missense alleles identified from WES. Twenty-seven cases did not undergo WES due to limitations in available material, but had known rare LoF or missense alleles identified by Sanger sequencing. Most putative causal mutations were typical LoF alleles or disrupted canonical mRNA splice sites (**Fig. 1C**). In agreement with previous reports, *RPS19, RPL5*, *RPS26*, and *RPL11* were the most frequently mutated RP genes (**Fig. 1B, E**). The majority of mutations were unique, with 80% of mutations observed in not more than one unrelated case (**Fig. 1D**). Sanger sequencing validated 100% of putative causal mutations identified. However, a small but considerable number of DBA gene mutations 7/472 (1.4%) were identified from targeted Sanger sequencing of the known DBA genes but were not found in the initial variant calls from WES (**Table S1**). While a few of these mutations were in genes duplicated in the hg19 genome build (*RPS17*), or in regions with low coverage (start site of *RPS24*), the majority were long and/ or complex indels. Thus, although WES is highly accurate, specific classes of clinically relevant LoF mutations, such as medium-sized indels, can be missed, and our results suggest a benefit to performing follow-up targeted Sanger or long-read sequencing when WES does not return a high-confidence causal mutation.

### Extended and Cryptic Splice Site Mutations in Known DBA Genes

Mutations that alter splicing, but that lie outside canonical splice donor or acceptor sites, including deep intronic variants, have recently been shown to account for a substantial fraction of Mendelian disease cases with previously unknown pathogenic variants^33; 48; 49^. Popular annotation tools, such as Variant Effect Predictor (VEP)^22^ and SnpEff^50^, define mutations that disrupt only the first two (GT/U) or last two (AG) intronic bases as canonical splice site mutations. Although WES can only detect mutations in sequences captured by exome baits and thus misses the majority of intronic bases, it can detect mutations in proximal splice sites. We observe not only an enrichment for canonical splice region mutations, but also a substantial increase in “extended” splice region mutations in known DBA genes in our cohort compared to 123,136 population controls from gnomAD (**Fig. S1**). While these mutations predominately affect the third base of the extended consensus splice acceptor or donor site (proband 1, **Fig. 2A, C**), we identified a small number of rare mutations further from the exon-intron junction that are not typically considered. For example, we identified a mutation 8 bases upstream of *RPS26* exon 3 (chr12:56437139:T>G) that was absent from gnomAD. This mutation is predicted to create a novel consensus acceptor site (TAT>TAG) that would likely result in a frameshift due to the inclusion of 7 additional nucleotides into the *RPS26* transcript (proband 3, **Fig. 2C**).

**Figure 2.**
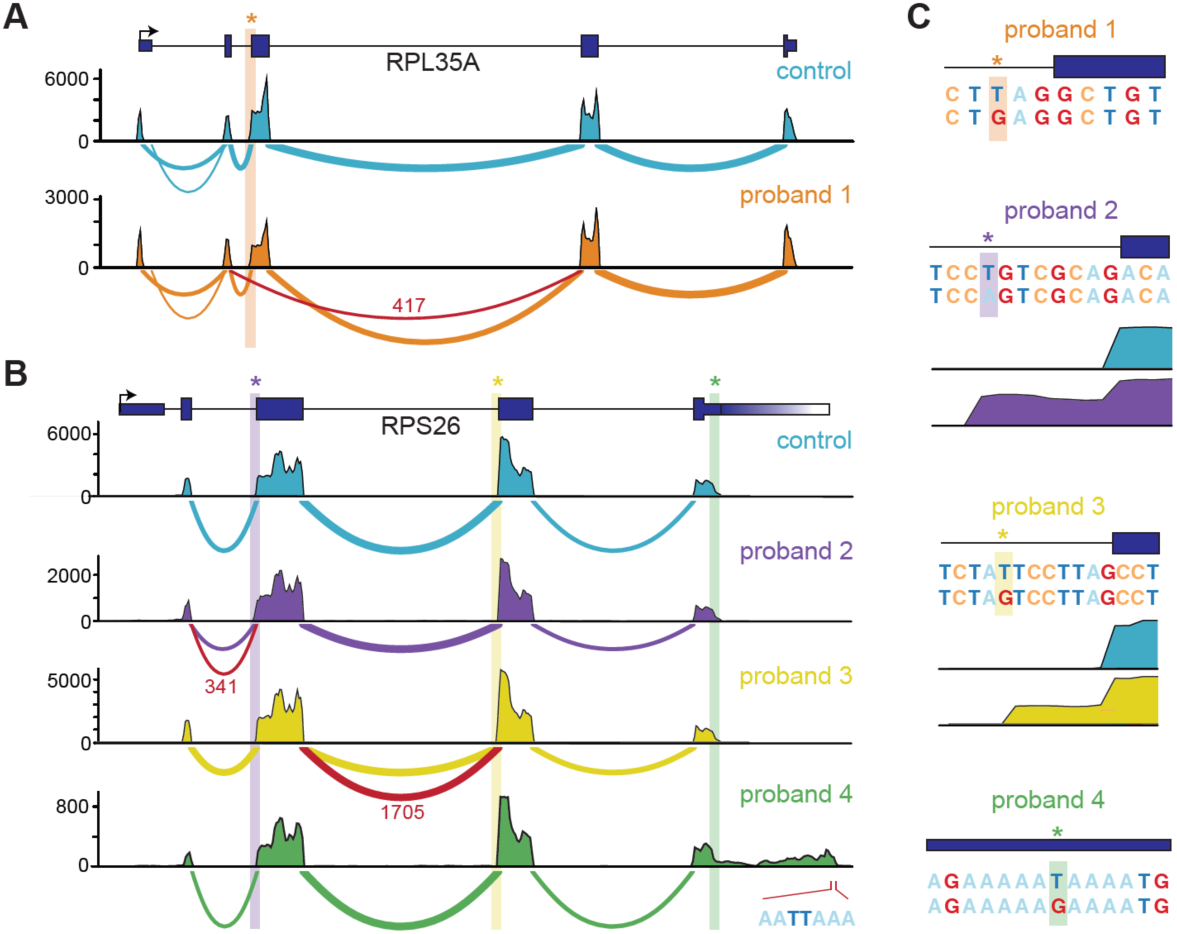
Non-Canonical Splice Variants in Known DBA Genes. (**A-B**) Sashimi plots of non-canonical splice mutants and a representative control are shown. The number of reads spanning each junction is indicated by the size of the sashimi plot curve, and the number of reads spanning novel junctions is indicated in red text. Coverage plots for total mapped reads are shown (total exon coverages is shown on the y-axis). Novel junctions due to the mutation indicated in (**C**) are highlighted in red. For the last panel in (**B**), the mutation disrupts a polyA binding site resulting in an extended 3’ UTR. The best guess novel polyA site used by this extended transcript is indicated. (**C**) Location and consequence of each mutation is shown, in addition to coverage plots for the exon extension mutants.

Since RP genes are ubiquitously expressed, we reasoned that performing RNA sequencing (RNA-seq) in cell lines derived from affected individuals would help us to determine if these extended splice region mutations were in fact splice-disrupting. Therefore, for 5 healthy controls and 9 cases with extended splice regions mutations, we created lymphoblastic cell lines (LCLs) and performed RNA-seq. For 6 of 9 cases, we observed aberrant splicing of the RP gene and/ or decreased mRNA expression (**Fig. 2A-C**, **Fig. S2**). In several cases, a mutation at the third base of a splice donor or acceptor site resulted in exon skipping (probands 1 and 5, **Fig. 2A-C**, **Fig. S2A**). Interestingly, for two unique *RPS26* mutations that were each 8 bases upstream of a different coding exon and created potential splice acceptor sites, we observed novel exon extensions (probands 2 and 3, **Fig. 2B-C**). In the case mentioned above, a novel acceptor site was created and used, resulting in a frameshift (proband 3, **Fig. 2C**). In another case, the presumed novel acceptor site was not faithfully used, again resulting in the introduction of a frameshift to the transcript (proband 2, **Fig. 2C**). The acceptor mutation in one individual was so severe that only limited splicing seemed to occur on the mutated transcript, and a substantial proportion of polyadenylated transcripts appeared to have intron retention (proband 7, **Fig. S2C-D**). Together, these results suggest that a proportion of cases lacking a typical RP gene mutation may instead harbor cryptic splicing mutations or mutations with posttranscriptional effects in one of the 19 currently known DBA genes. As the size of population-based WGS databases grows, identifying such cryptic mutations should become increasingly feasible with WGS. Furthermore, given the ubiquitous expression of RP genes, RNA-seq of patient-derived LCLs or fibroblasts could prove to be a relatively straightforward and fruitful strategy for identifying the functional impact on splicing or transcript expression from causal mutations missed by WES.

### Identification of a Null Mutation in the 3’ UTR of *RPS26*

We next extended our analysis to rare mutations that did not appear to be canonical LoF, missense, or splice region mutations in known RP genes. Interestingly, we identified a mutation in the 3’ UTR of *RPS26* that was also absent from gnomAD. This mutation was predicted to completely disrupt the polyadenylation signal (PAS) by changing the consensus motif AA(T/U)AAA to AAGAAA (proband 4, **Fig. 2C**). To test whether this was in fact the case, we created a LCL from the patient and performed RNA-seq. We found that transcription continued approximately 700 bases past the typical polyadenylation site, drastically increasing the size of the 3’ UTR (**Fig. 2B**). Furthermore, mRNA levels of *RPS26* were significantly reduced, although global mRNA profiles were largely similar (**Fig. S2E**). Although not tested here, it is likely that *RPS26* transcripts with the long mutant 3’ UTRs are less stable and targeted by miRNAs or RNA-binding proteins, resulting in reduced mRNA levels.

### Copy Number Variants in Known DBA Genes

In addition to missense or LoF mutations, smaller studies have estimated that 15-20% of DBA cases are due to a partial or full deletion of one copy of an RP gene^51; 52^. Although an imperfect approach, copy number variants (CNVs) can be identified as differences in coverage across regions ascertained by WES^36; 37^. Thus, we performed WES-based CNV calling (**Materials and Methods**) and identified 79 putative deletions in known DBA genes plus other RP genes (**Table S2**). To verify a subset of these deletions, we used digital droplet PCR (ddPCR) to test 13 of the most commonly deleted exons, representing 7 genes and 44 cases. Reflective of the fact that these putative deletions were carefully preselected as high confidence CNVs, 29 (66%) could be verified by ddPCR (**Fig. 3** and **Table S3**). *RPS17* was the most frequently deleted gene (11 cases, two verified as true *de novo*), predominately occurring as part of the well described 15q25.2 microdeletion^53^ (**Fig. 3D**). Notably, in hg19, *RPS17* is annotated as a duplicated gene (*RPS17* and *RPS17L*) and previous studies have considered whether a loss of one copy (out of 4) was sufficient to result in DBA^52^. Here, using empirical estimates of copy number from WGS for this region (**Fig. S3**), we determined that a duplication event is not supported and that there is in fact only a single copy of *RPS17* (indeed, this appears to have been resolved in hg38). Given our overall success here and in other studies of rare blood diseases^37^, we recommend that WES-based CNV calling should become a standard part of clinical WES analysis.

**Figure 3.**
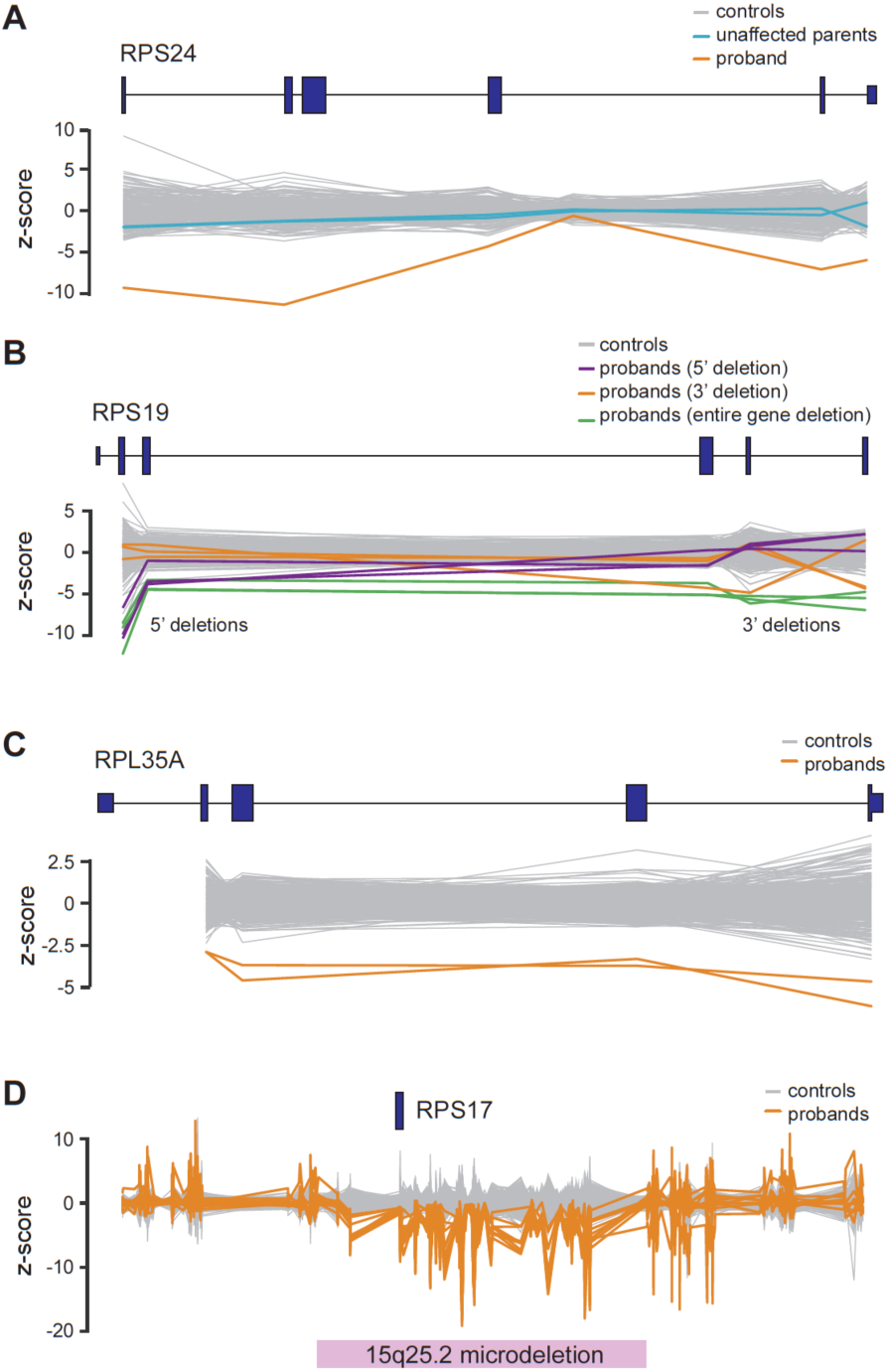
Copy Number Variants Identified by WES. Copy number variants identified by WES sequence are shown for (**A**) *RPS24*, (**B**) *RPS19*, (**C**) *RPL35A*, and (**D**) *RPS17*. A total of 558 controls consisting of other DBA samples or samples sequenced at the same time are shown in grey. (**A**) In this case, unaffected parents underwent WES and the inheritance of the deletion was determined to be *de novo*. (**B**) Several different deletions affecting only 5’ exons, 3’ exons, or the entire gene were detected across *RPS19*. (**D**) *RPS17* deletions were determined to almost exclusively be due to microdeletions of this region. A total of 11 individuals with *RPS17* deletions were detected.

### Prevalence and Penetrance of Mutations in Known DBA Genes

Although LoF and missense mutations occur far less frequently than expected in the majority of known DBA genes (**Table. S4**)^14^, the exact prevalence and penetrance of different allele frequency (AF) classes of DBA gene mutations have not been systematically investigated. We first investigated whether the class of more common but still rare (0.005% to 1%) missense mutations in DBA genes was enriched in cases compared to gnomAD, but observed a non-significant odds ratio of ≈ 1 (**Fig. S4A**). Since analyses of higher frequency variants may be confounded by unaccounted population structure, we used a set of unrelated dominant Mendelian genes as a control. We observed a larger enrichment for controls genes than for DBA genes, which suggests that, if anything, our results were biased against the null of no association between these variants and DBA (**Fig. S4A**; **Materials and Methods**).

There have been several reports of incomplete penetrance or variable expressivity for rare RP gene mutations^54-56^. Therefore, we set out to investigate the relative penetrance of rare DBA gene mutations^41^. Since no mutation was present in the DBA cohort at an allele count (AC) higher than 8 (14 including related individuals), we grouped mutations by both gene and type (LoF or missense). For the three most frequently mutated DBA genes (*RPS19*, *RPL5*, and *RPS26*), we found that LoF mutations demonstrate nearly complete penetrance (**Fig. 4A**). When we grouped LoF mutations in the less commonly mutated DBA genes, we similarly found that these mutations were also highly penetrant, although the point estimate was lower than for the more common DBA genes (**Fig. 4A**). This potentially suggests some variable expressivity of known DBA gene mutations, but this observation is confounded by the fact that not all predicted LoF mutations will cause a true loss of protein production. Since the majority of missense mutations were in *RPS19* (42/73, 58%), we investigated the penetrance of this class and found that rare missense mutations, in aggregate, were far less penetrant (6%) than rare LoF mutations (**Fig. 4A**). To determine if the missense mutations in the cohort were predicted to be more damaging than similarly rare missense mutations in gnomAD, we annotated all variants using Envision^57^, which, unlike most predictors^24; 47; 58; 59^, is not trained on gnomAD or databases of Mendelian mutations. Using Envision scores, we observed that damaging *RPS19* missense mutations were predicted to have higher penetrance (22%, **Fig. 4A**) and that the majority of RPS19 missense mutations in our DBA cohort were more damaging than those in gnomAD (**Fig. 4B**). However, we caution against over-interpretation, as not even predictive algorithms that were in part trained on RP gene mutations and/ or gnomAD could perfectly separate DBA missense mutations from gnomAD (auPRC range of 0.53 to 0.69).

**Figure 4.**
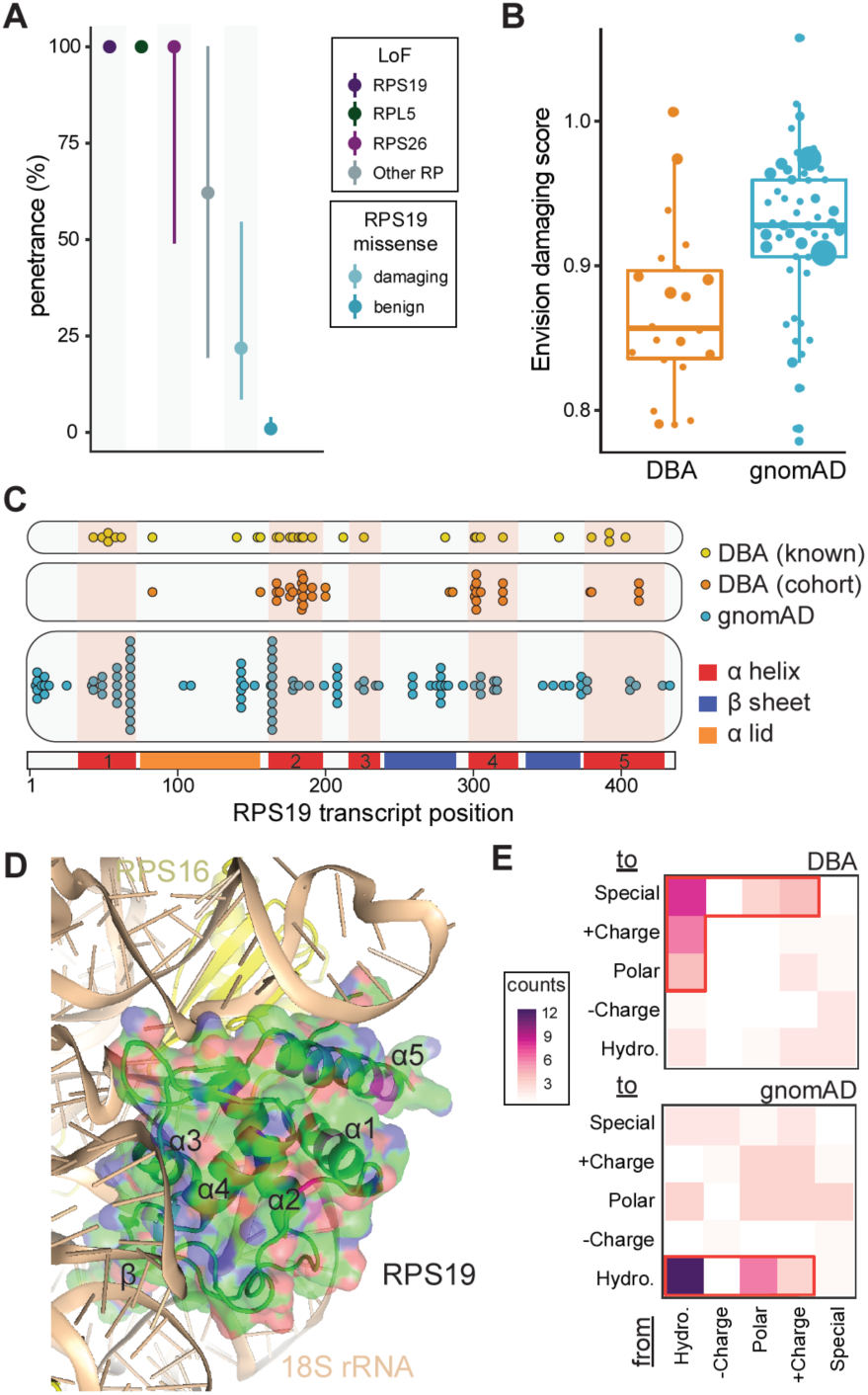
Penetrance and Prevalence of RP genes. (**A**) Near complete penetrance for LoF mutations in the top 3 most frequently mutated genes (57% of cases) was observed. Slightly lower estimates were obtained for LoF mutations in other known DBA-mutated RP genes. Penetrance was much lower for rare *RPS19* missense mutations (58% of all missense), but substantially increased when considering only predicted damaging mutations. (**B**) The majority of missense mutations identified in the DBA cohort are predicted to be damaging, whereas mutations of similar frequency in gnomAD are predicted to be benign. (**C**) *RPS19* missense mutations cluster into 3-4 groups along the mRNA transcript, although without clear separation from gnomAD mutations. (**D**) *RPS19* missense mutations appear to predominantly disrupt the stability of RPS19 by altering the hydrophobic core or by disrupting interactions with rRNA in the assembled ribosome (**Table S5**). The core α helices (1, 2, 4, and 5) and the β-hairpin are labeled. (**E**) Altering hydrophobic (Hydro.) amino acids to another type of amino acid was common in DBA but not in gnomAD. Special amino acids include Glycine, Proline, and Cysteine.

We next investigated the specific impacts of *RPS19* missense mutations, using both our cohort and a curated database of pathogenic DBA mutations (DBAgenes^60^). We observed three distinct types of mutations in our cohort. First, 48% of mutations changed a hydrophobic amino acid (AA) to a non-hydrophobic amino acid. Second, ≈ 18% of mutations changed a non-special AA to a special AA, such as proline. Third, 13% of mutations changed the smallest AA, glycine, to a much larger AA. These three types accounted for 78% of all DBA mutations but for only 25% of gnomAD mutations (p = 0.006). Structurally, we observed that 85% of *RPS19* missense mutation-carrying individuals in our cohort contained a mutation within exons encoding the four core α helices (1, 2, 4, and 5, **Fig. 4C**). Given that the locations of these mutations along the mRNA transcript were not fully independent from those variants observed in gnomAD, we investigated whether DBA mutations were more likely to affect specific structural elements of the RPS19 protein in the context of the fully assembled human ribosome (**Fig. 4D**). A high-resolution ribosome structure^42^ shows that 4 major α helices form a hydrophobic core to stabilize RPS19 (**Fig 4E**). Consistent with a previous report^61^ and our observation that DBA mutations often disrupt hydrophobic AAs, we determined that ≈ 50% of mutations would destabilize this hydrophobic core (**Table S5**). Furthermore RPS19 stabilizes two long hairpin ribosomal RNA (rRNA) regions at the head of the small subunit (**Fig. 4D**). These interactions would be disrupted by ≈ 43%, suggesting that the second largest class of missense mutations in RPS19 would disrupt interactions between RPS19 and rRNA, consistent with a previous hypothesis^61^.

Finally, motivated by a previous report that re-assessed the penetrance and pathogenicity of variants associated with Mendelian disease in public databases^41^, we investigated the frequency of reported variants from the DBAgenes database^60^ in the gnomAD population control. Importantly, 202/203 of the reported mutations in this database had 1 or fewer allele counts in gnomAD, consistent with our study as well as the low incidence and phenotypic severity of DBA. However, one missense variant (RPL5:c.418G>A, chr1:93301840:G>A) was relatively more common and was observed in 27 individuals in gnomAD, indicating that this variant was either not pathogenic or has low penetrance (**point estimate, (2.5-97.5% CI)**; 0.0%, 0.0-4.7%). Overall, our results indicate that rare LoF mutations in RP genes almost always result in DBA, whereas missense or more common mutations require increased scrutiny. Therefore, it is important for clinicians and researchers to rely on large population-based allele frequency estimates^46^, predictors of variant pathogenicity^24; 47; 57-59^, and other available clinical or experimental evidence before making a final determination of variant causality for DBA or similar disorders.

**Table 1.** Cohort Characteristics.

| DBA cases | no. | % |
| --- | --- | --- |
| Total | 472 |  |
| Families | 63 | 13.3% |
| Unrelated | 425 | 90.0% |
| WES (+ verification) | 445 | 94.3% |
| Sanger only | 27 | 5.7% |
| **Age at sample collection** |  |  |
| < 2 years | 138 | 32.1% |
| 2-10 years | 143 | 33.3% |
| 10-18 years | 57 | 13.3% |
| 18+ years | 92 | 21.4% |
| Unknown | 42 |  |
| **Sex** |  |  |
| Male | 249 | 53.4% |
| Female | 217 | 46.6% |
| Unknown | 6 |  |
| **Phenotype** |  |  |
| Typical | 269 | 85.9% |
| Mild | 15 | 4.8% |
| Atypical | 29 | 9.3% |
| Unknown | 159 |  |
| **Congenital malformations** |  |  |
| None | 159 | 55.6% |
| One or more | 127 | 44.4% |
| Head or face | 53 | 18.5% |
| Limbs | 44 | 15.4% |
| Genitourinary | 19 | 6.6% |
| Heart | 41 | 14.3% |
| Short stature | 25 | 8.7% |
| Unknown | 186 |  |
| **Elevated eADA** |  |  |
| Yes | 50 | 79.4% |
| No | 13 | 20.6% |
| Unknown | 409 |  |
| **Treatment** |  |  |
| Steroid dependent | 79 | 26.6% |
| No steroid trail yet | 49 | 16.5% |
| Transfusion dependent | 112 | 37.7% |
| Remission | 33 | 11.1% |
| BMT | 12 | 4.0% |
| No treatment | 12 | 4.0% |
| Unknown | 175 |  |

### Phenotype-Genotype Associations in Known DBA Genes

Although detailed phenotypic information was unavailable for a portion of the cohort (**Table 1**), we were nonetheless able to investigate phenotypic differences between individuals with disparate RP gene mutations (**Table S6**). This information was primarily obtained by report from referring clinicians or families. In agreement with previous studies on smaller cohorts (including a subset of this cohort)^2; 62; 63^, we observed significant differences in the presence of congenital malformations among individuals with mutations in different RP genes (**Fig. 5A**). In fact, the majority of individuals with *RPL5* (**point estimate, (2.5-97.5% CI)**; 83%, 67-93%) or *RPL11* (73%, 50-88%) mutations had one or more congenital malformations, in contrast to individuals with *RPS19* mutations where only 34% (24-47%) had any congenital malformation. In addition to observing significant associations between RP gene and congenital malformations affecting the head and face, limbs, stature, and genitourinary system, we also observed a significant association with the presence of congenital heart disease, which has previously been underappreciated^62; 64^ (**Fig. 5B**, **Fig. S5**). Leveraging the size of our cohort to make robust estimates, we conclude that 22% (12-35%) and 13% (4-31%) of individuals with *RPL5* and *RPL11* mutations, respectively, present with cardiac abnormalities, which is in stark contrast to the 4% (1-9%) and 7% (2-20%) of individuals with *RPS19* and *RPS26* mutations.

**Figure 5.**
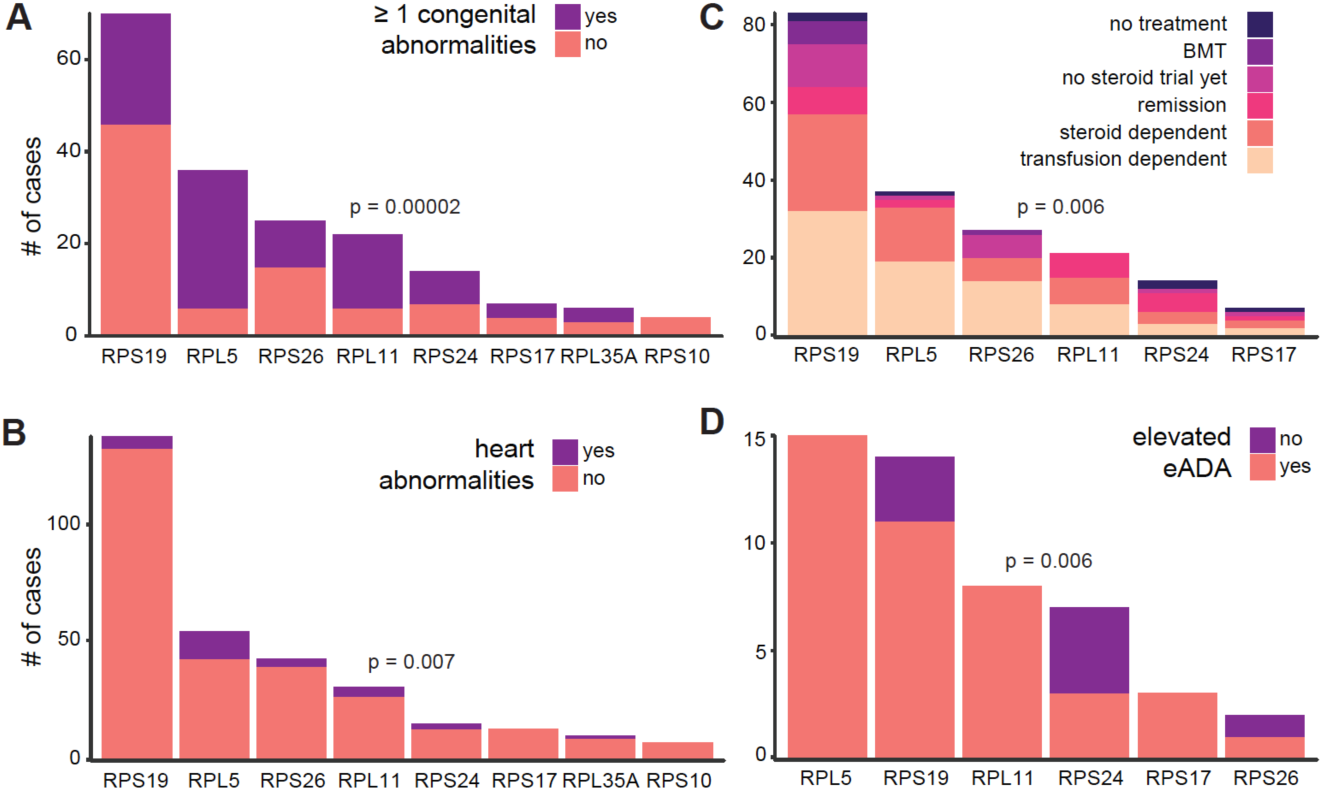
Phenotypic Associations. Differences in (**A**) presence of ≥ 1 congenital abnormality (**B**) heart malformations, (**C**) treatment requirements, and (**D**) eADA levels were observed between different RP genes. Differences remained significant after removing missense mutations. A Fisher’s exact test was used to test the hypothesis that there were differences in proportion of the outcome between RP genes. (**D**) The association between RP gene and treatment requirements appeared to primarily be due to differences in remission, and this association was no longer significant after removing individuals who went into remission

We next investigated whether there were differences in treatment requirements between RP genes for the primary condition of anemia, and observed a significant association (**Fig. 5C**). However, this did not appear to be due to differences in transfusion or corticosteroid treatment dependence, which account for approximately 65% of individuals (**Table 1**). Instead, we only observed a difference between RP genes for the proportion of individuals that were in remission (p = 0.001) (**Fig. S5E**). This appeared to be driven by the observation that 36% (14-64%) and 29% (12-52%) of individuals with *RPS24* and *RPL11* mutations were in remission and currently required no treatment, whereas only 8% (4-17%) and 5% (1-20%) of *RPS19* and *RPL5* individuals were in remission. After removing these individuals, the original association between RP gene and treatment requirement was no longer significant (p = 0.14), indicating that the major difference in treatment requirement between individuals with disparately mutated RP genes is the likelihood of remission.

Finally, we investigated whether there were differences in erythrocyte adenosine deaminase (eADA) levels, since elevated eADA is a useful diagnostic biomarker in DBA^65;66^. eADA measurement information was only available for 63 individuals and 79% were observed to have an elevated eADA, consistent with recent studies^65;66^. Although these studies reported little to no differences in eADA levels between RP genes, we observe a significant association where *RPS19* and *RPS24* individuals appear less likely to have elevated eADA (**Fig. 5D**). However, we caution that larger studies are required to determine if this observation is robust. Overall, these findings highlight the differences in clinical features due to disparate RP gene mutations.

### Novel Ribosome Protein Gene Mutations in DBA Cases

We next investigated whether we could identify additional RP genes involved in DBA. To identify putative causal mutations, we similarly searched for rare (gnomAD AC ≤ 3) LoF and missense mutations in the remaining 60 RP genes without previously reported DBA mutations. A total of 9 mutations (7 unique) involving 7 previously unreported RP genes were identified (**Table 2**). With the exception of *RPL10A*, the other 6 RP genes are extremely intolerant to mutation (pLI > 0.89), similar to nearly all other previously reported DBA genes (**Table. S4**)^14^. Two of the identified mutations were in splice regions and one altered a start codon. The other mutations were missense and were predicted by multiple algorithms to have damaging effects, whereas another mutation was in an RP gene encoded on the X-chromosome in a male individual. Although we do not explicitly validate any of these putative pathogenic mutations here, it is likely that many of these are novel RP genes causal for DBA (**Fig. 1E**). With further studies of independent cohorts of DBA patients, it is likely that additional evidence for a causal role of these genes may be established.

**Table 2.**
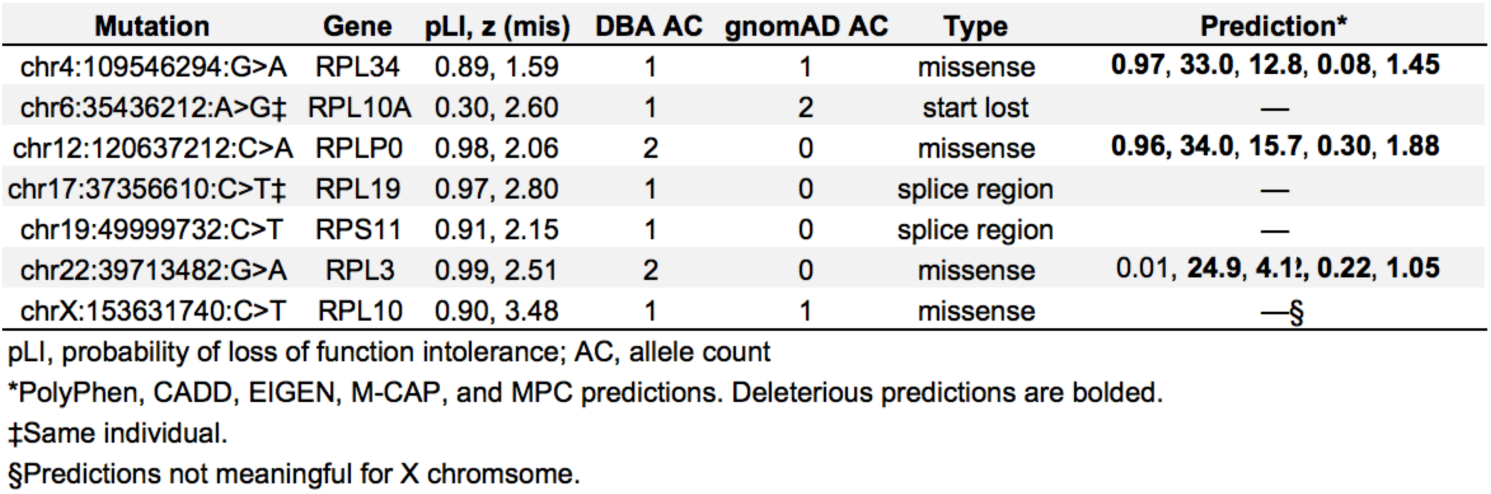
Novel RP Genes Mutated in DBA.

### Exome-Wide Significant Genes in DBA

Having extensively characterized mutations in the known DBA genes and other RP genes, we sought to identify novel genes associated with the clinical features characteristic of DBA by performing gene burden tests between unrelated individuals in our cohort and gnomAD controls (a cohort presumably depleted of rare pediatric diseases). We first carefully adjusted the variant quality thresholds between the cases and controls such that no genes were more enriched for rare (max DBA + gnomAD AC ≤ 3 or 6) synonymous mutations in the DBA cohort than expected (**Fig. 6A**; **Materials and Methods**). Restricting to rare LoF and damaging missense mutations with dominant inheritance, we identified *RPS19* (30% prevalence), *RPL5* (12%), *RPS26* (9%), *RPL11* (7%), and *RPS10* (1%) as significantly associated with DBA at an exome-wide significant threshold (p = 0.05 / 20,000) (**Fig. 6B-C**). If we additionally included all missense mutations, we observed a sub-threshold association for *RPL35A* (2%; p = 0.00001) with DBA. Together, mutations in these 5 genes account for 59% of DBA individuals in the cohort. However, we did not observe strong associations for the more prevalent genes *RPS24* (3%) and *RPS17* (3%), primarily because a large proportion of mutations in these genes were large deletions (we attempted to perform a CNV burden analysis, but were unable to properly control inflation). Among all other non-RP genes, we observed *SEH1L*, *HNRNPC*, and *ERCC1* as significantly associated with DBA in at least one test, but upon manual inspection the mutation calls were determined to be due to spurious alignments. Since we are both theoretically (**Fig. S6**) and empirically (**Fig. 6B-D**) well powered to detect genes containing mutations of clear effects, such as LoF or damaging missense we conclude that it is unlikely that dominant mutations in any single unknown gene that are detectable by WES are causal for more than 5% of cases of DBA with unknown genetic etiology.

**Figure 6.**
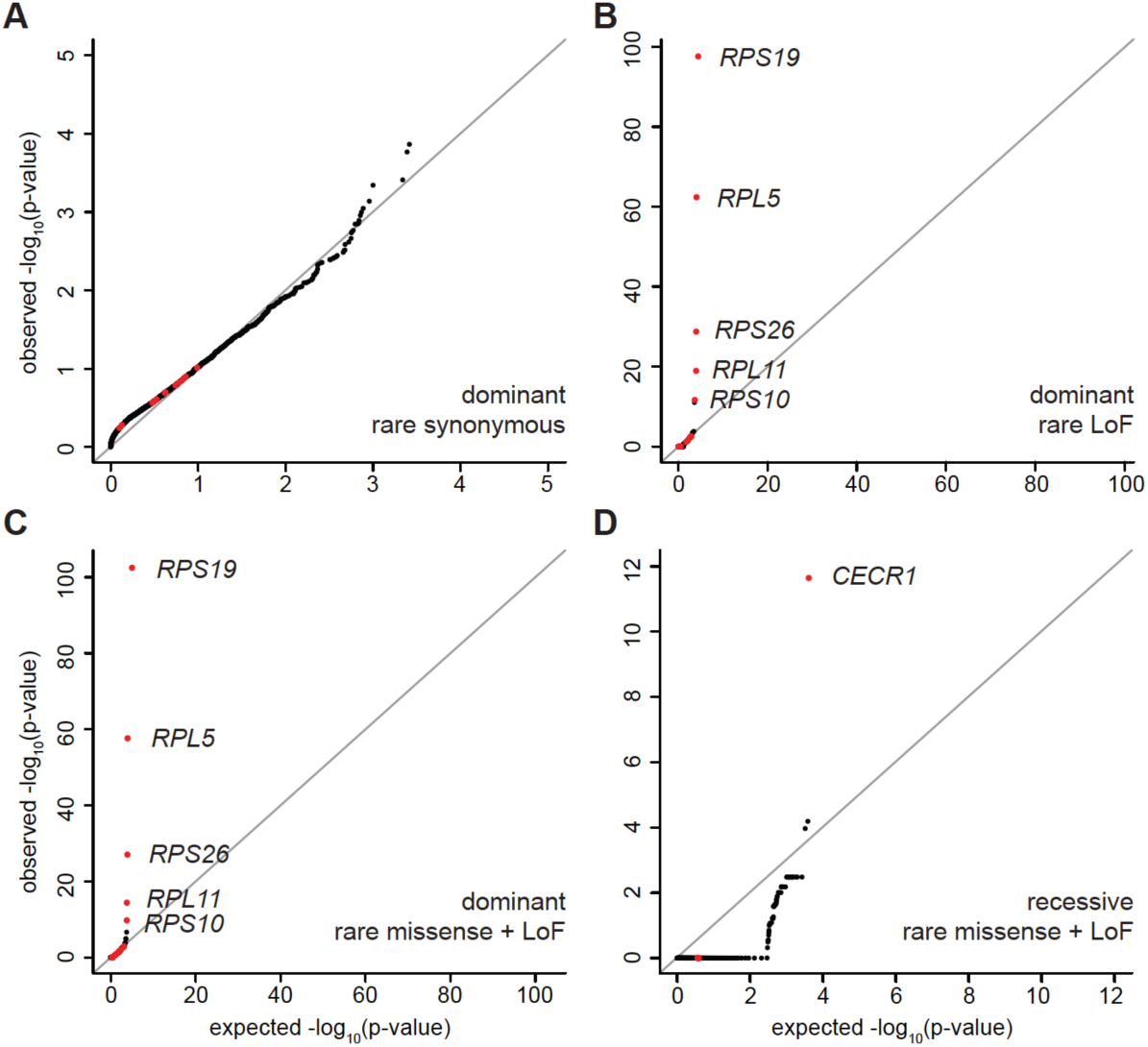
Gene Burden Results. (**A**) The burden of rare synonymous mutations in DBA indicate limited deviance from expected. (**B**) We observed an exome-wide significant association between rare LoF dominant mutations in *RPS19*, *RPL5*, *RPS26*, *RPL11*, and *RPS10*, 5 of the most commonly mutated DBA genes. (**C**) Similar results were observed after including rare damaging missense. (**D**) We observed an exome-wide association between rare LoF and damaging missense mutations with recessive inheritance for *CECR1* (*ADA2*). A fisher’s exact test was used to test for differences in each class of mutation between the DBA cohort and the gnomAD population control dataset, after filtering for high confidence variants and well covered regions consistent in both variant call sets.

Although we did not identify any new genes associated with DBA at the exomewide significant level for dominant inheritance, we performed similar gene burden tests for recessive inheritance. For rare (max DBA + gnomAD AC ≤ 20) LoF and damaging missense mutations with recessive inheritance, we identified one exome-wide significant gene, which was *CECR1* (**Fig. 6D**). In total, we identified 9 individuals with recessive or compound heterozygous missense or LoF mutations in *CECR1*, including 2 independent families in which recessive inheritance tracks with DBA status (**Table 3**). Although biallelic mutations in *CECR1* (that result in deficiency of ADA2; OMIM #607575) were initially associated with vasculitis, several recent reports have identified similar mutations in less than a handful of individuals diagnosed with DBA^67-69^. Preliminary evidence suggests that even though *ADA2* encodes an adenosine deaminase, similar to *ADA*, these individuals were not observed to have elevated eADA unlike the majority (85%) of DBA individuals. However, it is important to note that hematopoietic stem cell transplant appears curative in such individuals^67; 70^, suggesting that this disorder may emerge due to a hematopoietic intrinsic defect, although not necessarily intrinsic to the erythroid compartment itself. Overall, our data suggests that individuals presenting with DBA should be screened for *CECR1* mutations in addition to other known DBA genes and this is a condition that must be considered in any individual presenting with hypoplastic anemia.

### Phenocopies, Misdiagnoses, and non-RP Gene Mutations

Although *CECR1* was the only non-RP gene that was associated with a diagnosis of DBA at exome-wide significance, we investigated the extent to which there were other identifiable cases that either phenocopied or caused DBA. We conservatively identified 30 (6%) rare and predicted damaging genotypes in known or suspected red cell disorder genes that were non-RP genes (**Table 3**). Although the majority of these were in *CECR1* (9) or were mitochondrial deletions indicative of Pearson’s Syndrome (7), as has been previously reported^71^, we identified several genes of interest that were mutated in a small number of cases. First, our cohort contained five individuals with *GATA1* mutations^11; 16^, two related individuals with a shared *TSR2* mutation^13; 14^, and one individual with a rare *EPO* missense mutation^27^, each of which we have previously reported. We have shown that knockdown of TSR2, which is an RPS26 chaperone, mimics typical RP gene models of DBA^14^ and that altered GATA1 translation occurs due to RP haploinsufficiency^14; 16^, suggesting that these mutations result in DBA through common pathways. On the other hand, we observed that recombinant EPO containing the missense mutation altered EPO binding kinetics and that *in vivo* supplementation with wild-type EPO could rescue the hypoplastic anemia in a case with this mutation^27^. Since DBA is generally defined as a condition refractory to EPO treatment, the anemia caused by this mutation appears to represent a distinct clinical entity.

**Table 3.** non-RP Gene Mutations.

| Gene | Inheritance | # affected | Mutation 1 | Mutation 2 | Type | gnomAD AC |
| --- | --- | --- | --- | --- | --- | --- |
| Pearson (MT) | mitochondrial | 7 | mitochondrial deletion |  | deletion |  |
| CECR1 | recessive | 1 | chr22:17662818:A>T | chr22:17662818:A>T | missense | 0 |
| CECR1 | recessive | 1 | chr22:17662468:T>A | chr22:17662468:T>A | splice acceptor | 0 |
| CECR1 | recessive | 2 | chr22:17669238:C>T | chr22:17669238:C>T | missense | 6 |
| CECR1 | compound heterozygous | 1 | chr22:17690424:GC>G | chr22:17662785:T>C | frameshift, missense | 0, 2 |
| CECR1 | recessive | 3 | chr22:17662748:GTCAGCCT>G | chr22:17662748:GTCAGCCT>G | frameshift | 1 |
| CECR1 | recessive | 1 | chr22:17690423:GC>G | chr22:17690423:GC>G | frameshift | 0 |
| GATA1 | hemizygous | 2 | chrX:48649736:G>C |  | missense | 0 |
| GATA1 | hemizygous | 1 | chrX:48652176 C>T |  | splice region |  |
| GATA1 | hemizygous | 1 | chrX:48649735:AG>A |  | frameshift | 0 |
| GATA1 | hemizygous | 1 | chrX:48649518:T>C |  | start lost | 0 |
| SLC25A38 | recessive | 1 | chr3:39431108:G>T | chr3:39431108:G>T | splice donor | 0 |
| SLC25A38 | recessive | 1 | chr3:39436065:A>T | chr3:39436065:A>T | stop gained | 0 |
| TSR2 | hemizygous | 2 | chrX:54469851:A>G | chrX:54469851:A>G | missense | 0 |
| PUS1 | compound heterozygous | 1 | chr12:132414269:T>G | chr12:132426447:C>CT | stop gained, frameshift | 0, 0 |
| EPO | recessive | 1 | chr7:100320704:G>A | chr7:100320704:G>A | missense | 1 |
| NHEJ1 | recessive | 1 | chr2:220011458:G>A | chr2:220011458:G>A | stop gained | 2 |
| MYSM1 | recessive | 1 | chr1:59141211:G>A | chr1:59141211:G>A | stop gained | 0 |

AC, allele count

In addition to the previously reported DBA genes, we identified individuals with rare LoF variants in 4 genes that have been implicated in rare anemias that lacked a typical DBA gene mutation (**Table 3**). First, we identified two unrelated individuals with recessive LoF mutations (one novel, one known) in *SLC25A38*, a known causal gene for congenital sideroblastic anemia (CSA)^72^. Given the heterogeneous presentation of CSA, it is possible that the characteristic ring sideroblasts were initially limited or absent in samples obtained at diagnosis or that they were simply missed. In another individual, we identified compound heterozygous LoF mutations in *PUS1*, which encodes for pseudouridine synthase 1. Only a handful of individuals with *PUS1* mutations have been reported, but the majority presented with sideroblastic anemia, mitochondrial myopathy, and other dysmorphic features^73-75^. This individual died during infancy when the exact diagnosis may have been challenging to make. In another individual, we observed a novel recessive LoF mutation in *MYSM1*. Mutations in this gene have only been previously described in a handful of individuals^76; 77^, each presenting with transfusion-dependent refractory anemia in early childhood in addition to other cytopenias^78^. Finally, we identified one individual with a recessive LoF mutation in *NHEJ1*. Several individuals with similar mutations have been described as having immunodeficiency, dysmorphic faces, and in about half of cases, anemia and thrombocytopenia^79^. These findings suggest that DBA may be misdiagnosed in a small but important subset of individuals who in fact have one of a number of rare diseases in which hypoplastic anemia is a component of the phenotype.

## DISCUSSION

This work provides a systematic study on the approaches that can be used and the difficulties encountered when attempting to comprehensively define causal genetic lesions involved in a single Mendelian disorder. Even though we were conservative in assigning “causality”, we had a high genetic diagnosis rate of 78%, which is higher than other large reports on DBA cohorts and also higher than for most other Mendelian diseases. We achieved this high yet conservative diagnostic rate by leveraging large scale population genetic databases (gnomAD) to remove “common” variants down to an allele frequency of 0.003%, by using multiple, modern predictive algorithms when assigning pathogenicity, and by carefully investigating less well-annotated variants in or near known genes. To gain extra information from WES, we also used coverage information to nominate CNVs, 31 of which we orthogonally validated. Given the ubiquitous expression of RP gene mutations, we applied RNA-seq to LCLs derived from individuals with DBA and unambiguously validated 7 extended or cryptic splicing mutations, and a 3’ UTR mutation in *RPS26*. Finally, we did note a small, but significant, improvement in our genetic diagnostic rate by Sanger sequencing of known genes when WES was inconclusive, as Sanger sequencing can identify medium length and complex indels, a class of variants that is currently inadequately detected by WES approaches.

Although the phenotypic expression of DBA is largely homogenous, we observed that 6% of cases lacked typical mutations and instead harbored mutations that appeared to result in a phenocopy of DBA. Screening for causal mutations in DBA has typically been done using targeted Sanger sequencing of a handful of RP genes, but our work suggests that WES or WGS offers a substantial improvement. For example, we identified recessive *CECR1* mutations in 9 individuals in our cohort, highlighting the importance of screening for *CECR1* mutations in individuals with a clinical DBA diagnosis. Although we were well powered to identify novel genes harboring rare LoF or damaging missense alleles^45^, we did not identify any novel causal genes at an exome-wide significant level for DBA via gene burden testing. This suggests that larger sample sizes are needed to identify additional causal genes, even for rare, relatively homogeneous Mendelian diseases such as DBA. While we were able to carefully calibrate variant quality between gnomAD and our cohort, joint calling of genotypes would have likely improved our power as we erred on the side of being more conservative in our calibrations. Furthermore, as our ability to discriminate between benign and pathogenic variants improves, so will our ability to identify causal genes in Mendelian diseases. Assuming sufficient ascertainment of causal genetic variation, our results suggest that there is no single remaining gene with mutations detectable by WES that explains a large fraction (> 5%) of the remaining cases, as we would have almost certainly detected a burden of LoF or missense mutations in a gene of this character, given the cohort sample size. This leads us to believe that a large percentage of the remaining causal variants are RP gene CNVs, as previous studies have observed that 15-20% of cases harbor these, whereas our study detected only 10% since we did not use a comprehensive CNV screening assay. We also believe that we are only scratching the surface in identifying cryptic splice and large effect “non-coding” mutations (e.g. promoter, 3’ UTR, etc.) in RP genes. Comprehensively assaying CNVs will likely increase the diagnosis rate from 78% to 83-88% and combined WGS and RNA-seq on remaining cases could push the rate over 90%.

Overall, our results and other recent reports^80; 81^ suggest that at least 19 and perhaps 26 or more RP genes are involved in DBA pathogenesis (**Fig. 1E**). This is ≈ 1/3 of the genes that comprise the human ribosome, and mechanistic work from our group and others has suggested that these mutations predominately reduce ribosome levels, leading to a selective reduction in the translation of key genes involved in erythroid lineage commitment. However, there are still many unanswered questions. For example, it remains unclear if *CECR1* mutations result in an unrelated phenocopy or if CECR1 lies on the same causal pathway as other DBA mutations. It will also be interesting to examine to what extent other identified variants, such as the LoF mutations in *MYSM1*, may interface with the GATA1 pathway critical for erythropoiesis. Additionally, our work has built on other studies by demonstrating robust genotype-phenotype correlations, the detailed mechanisms of which remain to be elucidated.

## AUTHOR CONTRIBUTIONS

VGS, HTG, and JCU conceived of and designed the study. JCU, JMV, VGS, and HTG identified and interpreted putative causal mutations. JMV and JCU performed relatedness and ancestry analyses. LSL, DY, NA, and CF performed RNA-seq. JCU, JMV, and BBC performed RNA-seq and splicing analyses. JCU, REH, KRC, and ML performed CNV analyses. JMV and JCU performed penetrance analyses. AKK, JCU, JMV, and VGS performed and interpreted structural analyses. JCU performed phenotype-genotype associations. MHG, JCU, and RD performed gene burden analyses. RD, DGN, GG, EL, ML, AC, BBC, AHO-L, DGM, and SM provided assistance with analysis and interpretation. HTG, VGS, SK, AB, CG, CAS, PEN, EN, MM, AV, JL, EA, BG, AN, P-EG, M-FO, NM-L, and DJA compiled, phenotyped, and collected DNA for the cohort. NG, SBG, and ESL supervised DNA sequencing and variant calling. JCU, JMV, VGS, and HTG wrote the manuscript. All authors reviewed and edited the manuscript. VGS, HTG, and JCU supervised all aspects of this work.

## ACKNOWLEDGEMENTS

We owe a great deal of gratitude to the patients and families, without whom this study would not have been possible. We thank members of the Sankaran laboratory for valuable discussions and comments on this work. This work was supported by the National Institutes of Health grants R01 DK103794 and R33 HL120791 (to VGS), R01 HL107558 and K02 HL111156 (to HTG), UM1 HG008900 and R01 HG009141 (to DGM), as well as a DBA Foundation grant (to VGS). LDC, P-EG, and M-FO are supported by ANR (ANR 2015 AAP générique CE12-0001 - DBA Multigenes), and the EuroDBA project is funded by the ERA-NET programme E-RARE3 (ANR-15-RAR3-0007-04).

## SUPPLEMENTAL TABLE AND FIGURE LEGENDS

**Figure S1. Splicing Mutations at Known DBA Genes.** (**A**) Both canonical and extended splicing mutations are enriched for known DBA genes. No enrichment was observed for other RP genes or for other dominant Mendelian genes.

**Figure S2. Additional Non-Canonical Splice Variants in Known DBA Genes.** (**A-C**) Sashimi plots of non-canonical splice mutants and a representative control are shown. The number of reads spanning each junction is indicated by the size of the sashimi plot curve. Novel junctions due to the mutation indicated in (**A-B**) are highlighted in red. (**D**) Location and consequence of each mutation is shown, in addition to coverage plots for the exon extension mutant (**B**) and intron retention mutant (**C**). (**E**) Log^2^ fold change in transcripts per million across annotated RP genes for the indicated proband vs. 5 control LCLs.

**Figure S3. Re-evaluations of *RPS17* Copy Numbers using WGS.** (**A**) Using 2,535 individual samples that underwent WGS from the 1000 Genomes Project, we estimated the copy number of the annotated segmental duplication containing *RPS17* (and *RPS17L*) in hg19. Nearly all samples have coverage indicative of only 2 copies for the *RPS17* “segmental duplication”, rather than 4 copies, as can be observed in the control segmental duplication of approximately the same size.

**Figure S4. No Evidence for More Common DBA Mutations.** (**A**) No enrichment is observed for more common mutations in known DBA genes. There is a small but significant enrichment observed for other dominant Mendelian genes, indicating a possible mismatch unmodeled confounding in variant filtering or in population stratification. (**B**) Results from 6 well known missense variant effect predictors indicate that DBA *RPS19* mutations are more damaging than gnomAD *RPS19* mutations. However, no predictor can perfectly separate the two groups.

**Figure S5. Additional Phenotypic Associations.** Differences in (**A**) head or craniofacial abnormalities, (**B**) limb or hand abnormalities, (**C**) genitourinary abnormalities, (**D**) short stature or skeletal abnormalities, and (**E**) remission status were observed between different RP genes. A χ^2^ test was used to test the hypothesis that there were differences in proportion of the outcome between RP genes.

**Figure S6. Gene Burden Power Analysis.** (**A**) Power for different allele count scenarios was calculated using a 1-sided Fisher’s exact test. Within the quality normalized gnomAD dataset, we observe that the top 10 and 25% of constrained genes have fewer than 4 or 10 rare LoF allele counts in gnomAD, respectively. Thus, we consider genes with 0-10 rare LoF as “constrained” genes, similar to the RP genes already implicated in DBA (median of 0 rare LoF alleles in gnomAD). In this scenario, the gene burden tests were well powered (> 80%) to detect an exome-wide significant gene association with as few as 4 to 6 individuals with LoF mutations (corresponding to 1-1.5% of DBA incidence). Similarly, we observe that the top 25% of constrained genes have 165 missense mutations and 53 predicted damaging missense mutations (median of 59 and 8 for RP genes implicated in DBA, respectively). Thus, we consider 100 counts in gnomAD as a reasonable number of rare missense alleles for a “constrained” gene, but we also consider an extreme scenario of up to 1,000 allele counts. In both scenarios, gene burden tests were well powered (> 80%) to detect an exome-wide significant gene association with as few as 8 to 16 individuals with missense mutations (corresponding to 2-4% of DBA incidence). Thus, we are theoretically well powered to detect mutations between 1-4% of total DBA incidence. However, after conservatively adjusting the variant quality threshold in our DBA cohort specifically for burden analysis, we were unable to detect exome-wide significant associations for RPL35A (2%) and RPS24 (3%) as several validated variants were filtered due to lower quality scores in WES, but we could detect exome-wide significant associations for RPS10 (1%) and RPL11 (7%). Given these considerations, we believe that a more appropriate, if slightly conservative, estimate of our true power lies closer to ≈ 5%.

**Table S1. Variants Missed By WES But Identified By Sanger Sequencing.**

**Table S2. Putative CNVs Identified by WES.**

**Table S3. WES-based CNV Validation.**

**Table S4. DBA Genes Are De-enriched For LoF And Missense Variants.**

**Table S5. Predicted Structural Impacts of *RPS19* Missense Mutations.**

